# Safety switch optimization enhances antibody-mediated elimination of CAR T cells

**DOI:** 10.1101/2022.08.24.505164

**Authors:** Tamer B Shabaneh, Howell F Moffett, Sylvia M Stull, Thomas Derezes, Leah J Tait, Spencer Park, Stan R Riddell, Marc J Lajoie

## Abstract

Activation of a conditional safety switch has the potential to reverse serious toxicities arising from the administration of engineered cellular therapies, including chimeric antigen receptor (CAR) T cells. The functionally inert, non-immunogenic cell surface marker derived from human epidermal growth factor receptor (EGFRt) is a promising safety switch that has been used in multiple clinical constructs and can be targeted by cetuximab, a clinically available monoclonal antibody. However, this approach requires high and persistent cell surface expression of EGFRt to ensure that antibody mediated depletion of engineered cells is rapid and complete. Here we show that incorporating a short juxtamembrane sequence into the EGFRt polypeptide enhances its expression on the surface of T cells and their susceptibility to antibody-dependent cellular cytotoxicity (ADCC). Incorporating this optimized variant (EGFRopt) into bicistronic and tricistronic CAR designs results in more rapid *in vivo* elimination of CAR T cells and robust termination of their effector activity compared to EGFRt. These studies establish EGFRopt as a superior safety switch for the development of next-generation cell-based therapeutics.

## INTRODUCTION

The clinical success of CAR T cells targeting lineage-restricted antigens expressed in B cell leukemias and lymphomas is tempered by toxicities in a subset of patients.^1^ Many target molecules being pursued in other malignancies are also expressed on nonexpendable normal tissues, increasing the potential for severe on-target off-tumor toxicities. Indeed, serious toxicity has been reported in CAR T cell trials in both hematologic and solid cancers and attributed to recognition of normal cells.^2–4^ Insertion of CAR constructs into T cells also carries a risk of insertional mutagenesis and oncogenic transformation.^5,6^ These and other toxicities are potentially reversible if a conditional safety switch is incorporated such that, when targeted, it would swiftly and selectively eliminate adoptively transferred T cells in patients.

Several safety switches have been developed with this goal. Such strategies include the co-expression of proteins that respond to small molecule drugs, including inducible caspase-9 (iCasp9),^7^ herpes simplex virus tyrosine kinase (HSV-TK)^8^, or human thymidylate kinase (TMPK).^9^ However, the immunogenicity of virally derived HSV-TK can result in the unwanted elimination of therapeutic cells when toxicity is absent.^10,11^ High expression levels of HSV-TK are necessary for efficient elimination of engineered cells and several days may be required for elimination.^12^ By contrast, iCasp9, which is human derived and less likely to be immunogenic, can be rapidly dimerized by a small molecule drug resulting in apoptosis of cells with high expression levels of the transgene.^13^ In settings of hematopoietic stem-cell transplantation, iCasp9-expressing donor T cells can be rapidly eliminated to reverse organ toxicity resulting from graft-versus-host disease (GVHD) mediated by T cell recognition of host alloantigens.^14^ However, the small molecule dimerizer is not available as an FDA-approved drug, and the efficiency of iCasp9-mediated elimination when the transgene is expressed in a multicistronic construct, such as in CAR T cells, requires clinical evaluation.

In contrast to small-molecule targeting, expression of cell-surface markers on T cells facilitates monitoring by flow cytometry (for manufacturing and for pharmacokinetics) and enables targeted elimination by administering target-specific antibodies. Such approaches include human epidermal growth factor receptor (EGFR),^15^ RQR8,^16^ or CD20,^17^ and allow for the use of clinically approved antibodies, such as cetuximab or rituximab, to target infused populations for elimination. EGFR is a receptor tyrosine kinase expressed on many cancer cells and can be targeted with cetuximab.^18^ Removal of the EGFR N-terminal extracellular ligand-binding domains I and II along with the intracellular signaling modules preserves the cetuximab epitope and retains cell surface expression. This truncated EGFR (EGFRt) was developed as a transduction marker and safety switch for clinical T cell therapy^15,19–22^ and demonstrated to function as a target for antibody-mediated elimination in preclinical models.^23^ However, as cetuximab-mediated antibody-dependent cellular cytotoxicity (ADCC) correlates with EGFR expression levels, cetuximab elicits slow and incomplete ablation of CAR T cells with low-EGFR expression.^23,24^ Therefore, to maximize the utility of truncated EGFR for *in vivo* depletion, we sought to increase the surface expression and stability of the marker. We demonstrate that incorporating a short juxtamembrane sequence into a truncated human EGFR-derived polypeptide increases its cell surface expression, leading to markedly enhanced cetuximab-mediated *in vitro* ADCC and *in vivo* CAR T cell ablation compared to the previously described EGFRt. The optimized truncated EGFR variant (EGFRopt) can be used as a superior safety switch to facilitate development of next-generation CAR T cell therapeutics.

## RESULTS

### Basic or glycine-rich juxtamembrane sequences substantially increase surface expression of EGFR-derived proteins

Receptor tyrosine kinases, such as EGFR, share a common architecture consisting of an extracellular ligand binding domain, a single-pass transmembrane helix followed by a flexible juxtamembrane domain, a protein kinase, and a C-terminal region (Figure 1A). Positively charged residues can drive transmembrane domain topology^25^ and contribute to the free energy of transmembrane domain insertion.^26^ Consistent with this, the juxtamembrane region of human EGFR is enriched in basic residues, which can form a structured interaction with the membrane^27^ (Figure 1A-B). To test the impact of juxtamembrane sequences on the surface display of truncated EGFR molecules, we generated modules containing a synthetic GMCSF-derived signal peptide, EGFR domains III-IV, EGFR transmembrane domain, and short intracellular domains derived from the native EGFR sequence or synthetic sequences (Figure 1B). Since T678 of human EGFR may be a site of regulatory phosphorylation, we limited the tested sequences to amino acids 669-677 (RRRHIVRKR). Furthermore, glycine residues can disrupt α-helical structure, exposing the peptide backbone, so we also tested intracellular domains containing unstructured glycine/serine-rich linkers (S-G4Sx2), or a short portion of the native juxtamembrane domain followed by a glycine/serine-rich linker (RRRS-G4Sx2).

**Figure 1:**
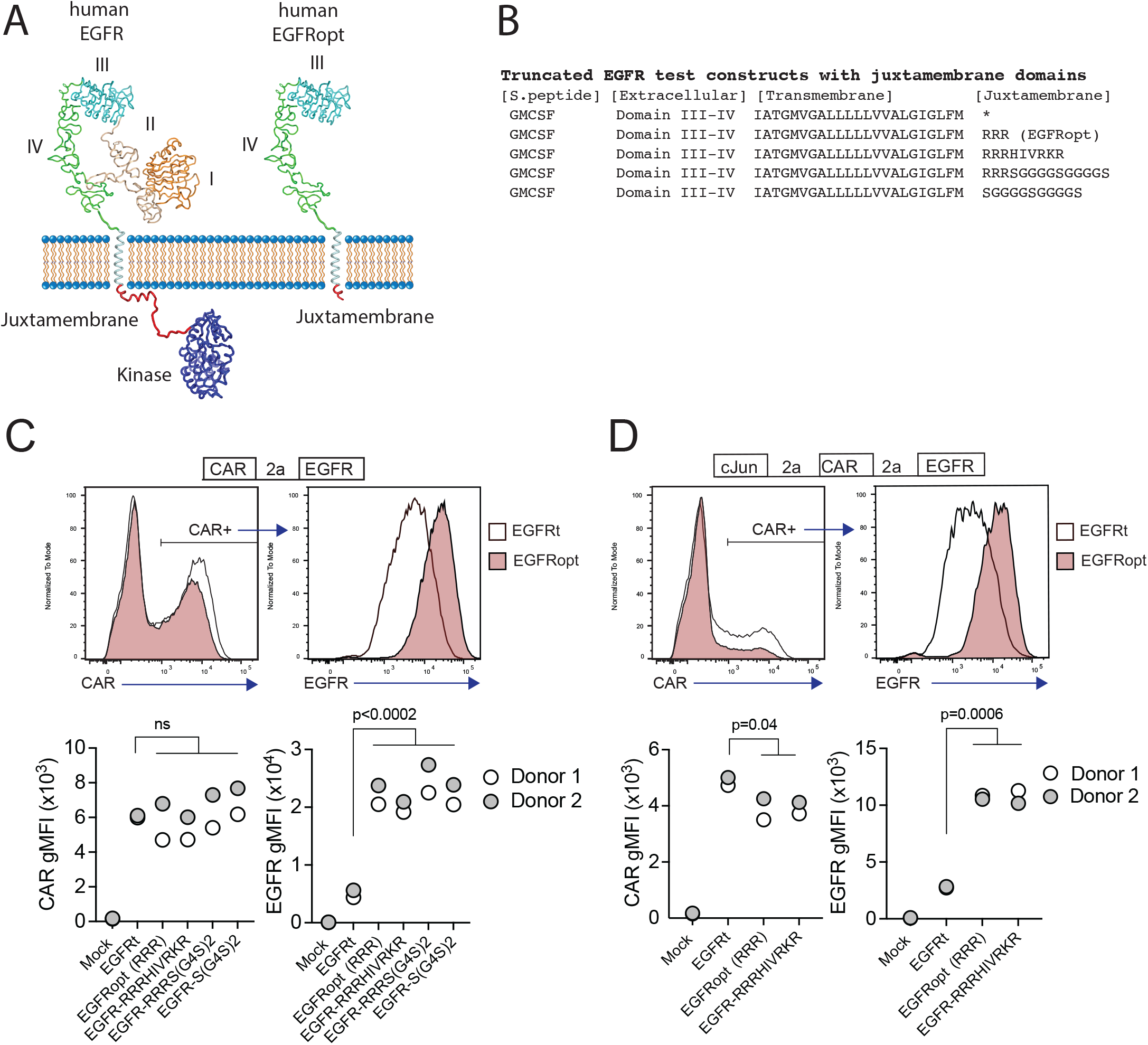
Basic or glycine-rich juxtamembrane sequences strongly increase surface display of EGFR-derived proteins. (A) Structural depiction of human EGFR (left), and EGFRopt containing a short juxtamembrane anchor sequence (right) based on crystal structures of EGFR extracellular domains I-IV (PDB ID 1YY9^44^), kinase domain (PDB ID 3GOP^45^), and nuclear magnetic resonance structures of EGFR transmembrane domain and juxtamembrane domains in lipid micelles (PDB ID 2N5S^27^). Protein sequence C-terminal of the kinase domain is not shown. (B) Peptide sequence of truncated EGFR protein modules including juxtamembrane anchor sequences. (C-D) Expression of truncated EGFR modules from bicistronic (C) and tricistronic (D) CAR constructs gated on CAR+ cells. Top: representative histograms of surface detection of CAR and EGFR on cells transduced with EGFRt and EGFRopt; Bottom shows gMFI of CAR and EGFR for various test sequences in 2 donors. Symbols represent 2 donors; analyzed by ANOVA (Dunnet post hoc).

Primary T cells were transduced with lentiviruses encoding a ROR1-specific CAR linked by a P2A skip sequence to the various EGFR variants. The co-expression of the CAR and EGFR-derived polypeptides were measured by flow cytometry. EGFR variants that included a basic or glycine-rich juxtamembrane sequence consistently demonstrated higher mean fluorescence intensity, as compared to the previously described EGFRt, whereas surface expression of the translationally linked ROR1 CAR was consistent for all constructs (Figure 1C). The relatively higher expression of EGFR-derived polypeptides was observed on T cells transduced over a wide range of viral MOIs and was optimal when at least two juxtamembrane arginine residues were used (Figure S1A-B). These data demonstrate that the addition of certain juxtamembrane sequences markedly increases the level of truncated EGFR stably expressed on the T cell surface.

CAR constructs often incorporate additional transgenes—such as the transcription factor c-Jun—to augment therapeutic efficacy.^28^ However, P2A peptide linkage can negatively affect expression of genes towards the terminal end of multi-gene cassettes.^29^ To determine if juxtamembrane sequences maximizes the surface expression of EGFR variants in tricistronic formats, we evaluated constructs that encoded c-Jun, ROR1 CAR, and truncated EGFR variants that include no juxtamembrane sequence (EGFRt), three arginine residues (referred to henceforth as EGFRopt), or nine juxtamembrane residues (EGFR-RRRHIVRKR) (Figure 1D). As observed with bicistronic vectors, the surface expression of EGFR variants having juxtamembrane sequences increased by about four-fold compared to EGFRt, while minimal reduction in CAR expression was observed. Collectively, these studies indicate that the inclusion of a juxtamembrane domain boosts the cell surface expression of truncated EGFR, and that the 3 amino acid RRR sequence was sufficient for this effect.

### EGFRopt enhances cetuximab-mediated lysis of CAR T cells *in vitro*

Cetuximab mediates antibody-dependent cellular cytotoxicity (ADCC) of target cells and activity is dependent on the level of surface EGFR expression.^24^ We hypothesized that higher surface expression of truncated EGFR would improve its utility as a safety switch in CAR T cell therapy through enhanced ADCC of CAR-transduced cells. As the four variant juxtamembrane sequences achieved similar increases in surface expression of EGFR, these studies focused on comparing EGFRt (no juxtamembrane sequence) with EGFRopt (RRR juxtamembrane sequence) (Figure 2A). CAR T cells expressing EGFRt were only modestly susceptible to ADCC following 4-hour treatment with NK cells and cetuximab, whereas T cells expressing EGFRopt were efficiently targeted for NK-mediated ADCC, even at lower doses of cetuximab (Figure 2B).

**Figure 2:**
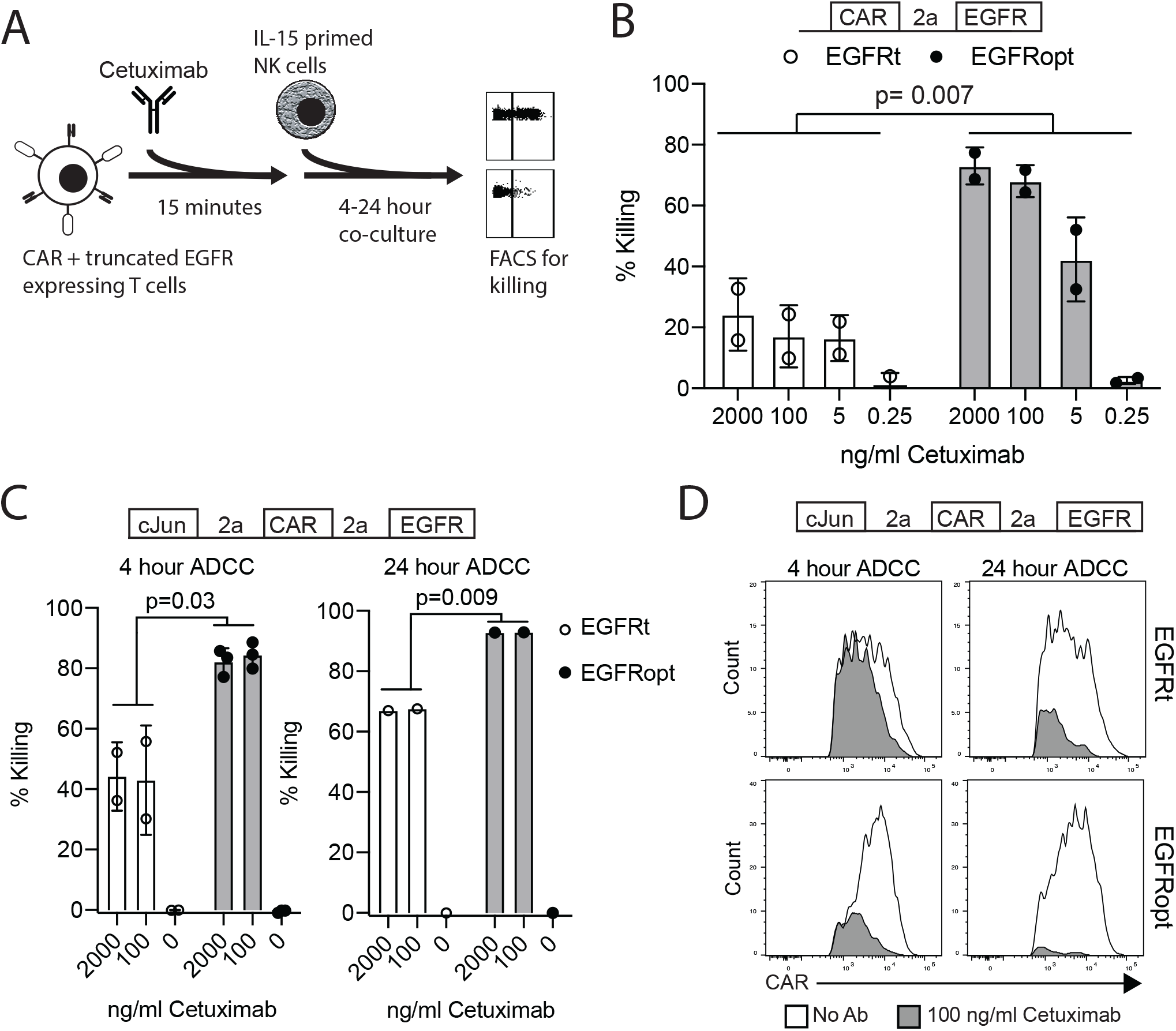
EGFRopt enhances the targeting of CAR T cells for ADCC *in vitro*. (A) Schematic of the ADCC protocol: CAR-T cells were pre-incubated with cetuximab biosimilar before co-culture with IL-15 primed NK cells. After co-culture, ADCC was assessed by flow cytometry for residual CAR-positive T cells. (B) Cetuximab-mediated ADCC of T cells transduced with bicistronic constructs after 4 hrs co-culture. (C) Cetuximab-mediated ADCC of T cells transduced with tricistronic constructs after 4 or 24 hours. (D) Representative flow plots showing relative CAR expression on remaining CAR+ cells from C in the presence (gray) or absence (white) of cetuximab. Symbols in B and C represent biological replicates. Symbols represent 2-3 donors; analyzed by ratio paired t test.

In the tricistronic setting where the EGFR marker expression is lower, minimal ADCC was observed when targeted T cells expressed EGFRt (Figure 2C, left), whereas expression of EGFRopt markedly improved sensitivity to cetuximab-mediated ADCC (Figure 2C, right). Increasing co-culture duration improved ADCC-mediated T cell ablation, with near complete elimination of EGFRopt T cells compared with ~70% killing of EGFRt cells (Figure 2C, right). ADCC of EGFRt CAR T cells was largely restricted to the highest CAR expressing cells at both time points, whereas only a minor population of very low CAR expressing EGFRopt T cells remained viable (Figure 2D). These data indicate that addition of juxtamembrane domains to truncated EGFR enhances its safety-switch function, even in settings involving low transgene expression.

### EGFRopt enhances cetuximab mediated clearance of CAR T cells *in vivo*

Paszkiewicz *et al.* previously showed that murine CAR T cells expressing EGFRt can be eliminated *in vivo* by administering cetuximab.^23^ Murine CD8 T cells transduced with EGFRopt exhibited more robust surface expression compared to EGFRt, despite similar T cell-transduction efficiencies (Supp Fig S1C-D). To determine whether cetuximab eliminated EGFRopt CAR T cells more rapidly than EGFRt CAR T cells *in vivo*, we adoptively transferred murine-CD19-targeting CAR T cells (m19.CAR T) and monitored their levels in blood 3 hours following treatment with cetuximab (Figure 3A-B). EGFR mRNA transcript levels were used as surrogate for CAR T cell levels due to the high sensitivity of qRT-PCR. Mice engrafted with EGFRopt CAR T cells exhibited a sharper drop (44-fold) in EGFR transcript levels post cetuximab compared to mice engrafted with EGFRt CAR T cells (6-fold). The depletion kinetics of CAR expressing T cells were confirmed by generating a m19.CAR containing a STII tag in the hinge region to allow direct staining of CAR T cells *in vivo*^30^. As expected, a sharper drop in the frequency of STII+ EGFRopt CAR T cells compared to EGFRt CAR T cells was observed 3 hours post cetuximab (14-fold vs. 7-fold) (Figure 3C). Notably, frequencies of EGFRopt CAR T cells in the bone marrow 24 hours after cetuximab treatment were significantly decreased compared to EGFRt CAR T cells (9-fold vs 2-fold), underscoring the efficacy of EGFRopt as a rapid safety switch (Figure 3D). Therefore, EGFRopt facilitates more rapid and profound cetuximab-mediated depletion of CAR T cells *in vivo*.

**Figure 3:**
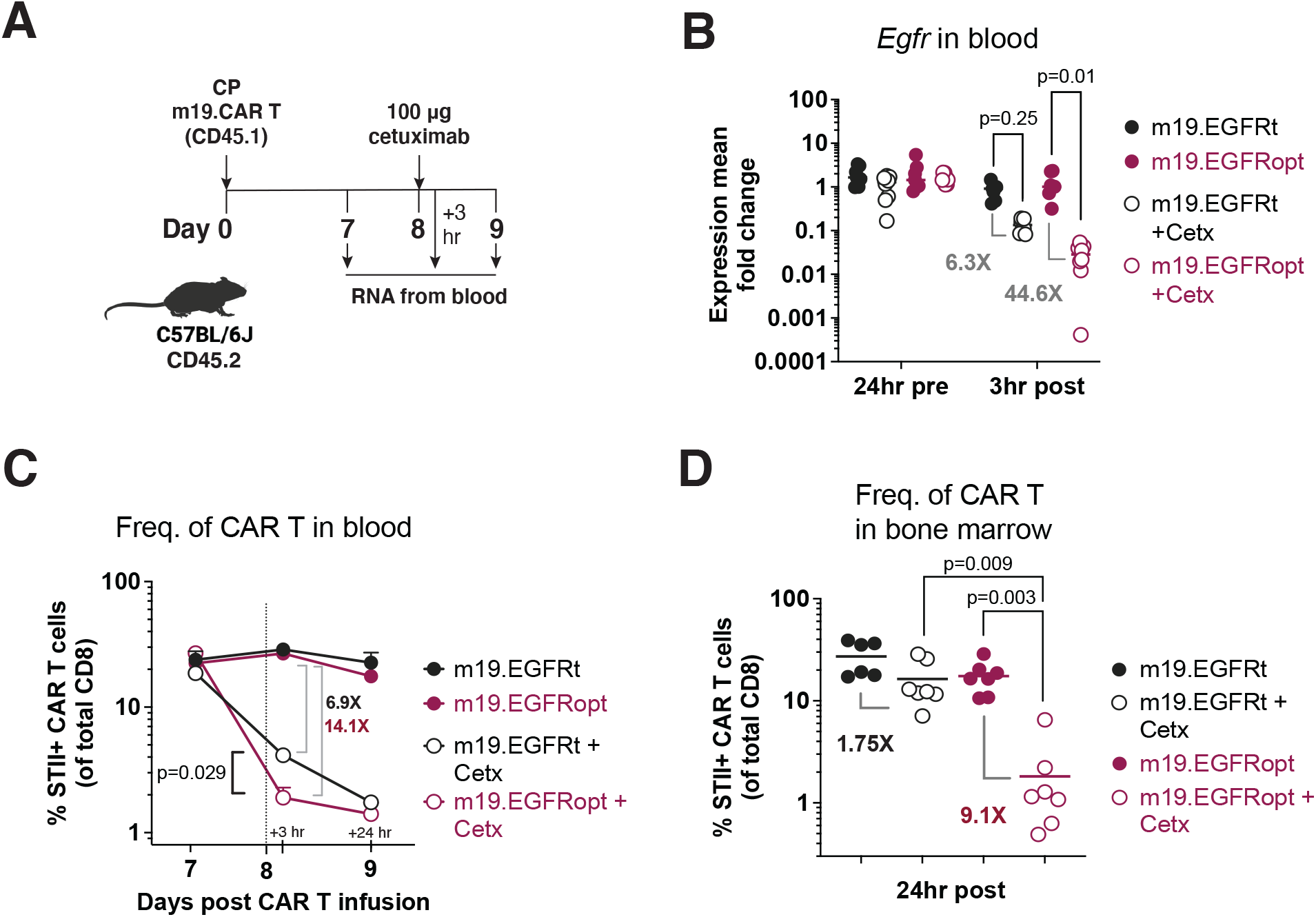
EGFRopt enhances antibody mediated depletion of CAR T cells *in vivo*. (A) Schematic of the CAR T cell-depletion experiment in B-D. Mice were lymphodepleted with cyclophosphamide (CP) and infused with EGFRt or EGFRopt-marked CAR T cells. A week later, mice were treated with 100 μg cetuximab or left untreated. (B) Quantitative RT-PCR analysis of *egfr* levels in cetuximab-treated vs. untreated mice. (C) CAR T cell frequencies in blood quantified by flow cytometry 3 hours and 24 hours post cetuximab, represented by the vertical line. (D) Frequency of CAR T cells in bone marrow 24 hours post cetuximab. (B) n=7-8, or (C-D) n=6-7 mice/group; symbols and error represent means ± SEM. Absence of error bar indicates SEM less than area represented by symbol. Analyzed by ANOVA (Šídák post hoc).

### Superior elimination of CAR T cell effector activity with EGFRopt promotes rapid recovery of normal B cells

Cetuximab administration resulted in more profound depletion of m19.CAR EGFRopt T cells, which we hypothesized would result in more rapid recovery of circulating CD19+ B cells. m19.CAR T cells expressing EGFRt or EGFRopt were adoptively transferred into lymphodepleted mice, and T cell engraftment and B cell aplasia were monitored in the blood over time (Figure 4A). As expected, a single cetuximab dose depleted EGFRt and EGFRopt CAR T cells from blood, and CAR T cell levels remained suppressed throughout the observation period (Figure 4B). Mice administered m19.CAR T cells without cetuximab exhibited a decline in the frequency of CAR T cells as lymphocyte counts recovered from lymphodepletion but had sustained B cell aplasia (Figure 4B, C). By contrast, cetuximab-treated mice exhibited B cell recovery within 2 weeks of drug administration in the EGFRopt cohort, which was a week earlier than the EGFRt cohort (Figure 4C). Cohorts of mice were also treated with tricistronic CAR T cells with the same outline in Figure 4A, and B cell aplasia was tracked in the blood over time (Figure 4D). Mice treated with c-Jun.m19 CAR T cells exhibited sustained B cell aplasia, whereas B cells in cetuximab-treated mice began to rebound on Day 10 post depletion in the EGFRopt cohort, 18 days earlier than the EGFRt cohort (Figure 4E, F). Collectively these data show that EGFRopt exhibits superior function as a safety switch compared to EGFRt, and results in more rapid recovery of normal CD19+ B cells from hematopoietic progenitor cells.

**Figure 4:**
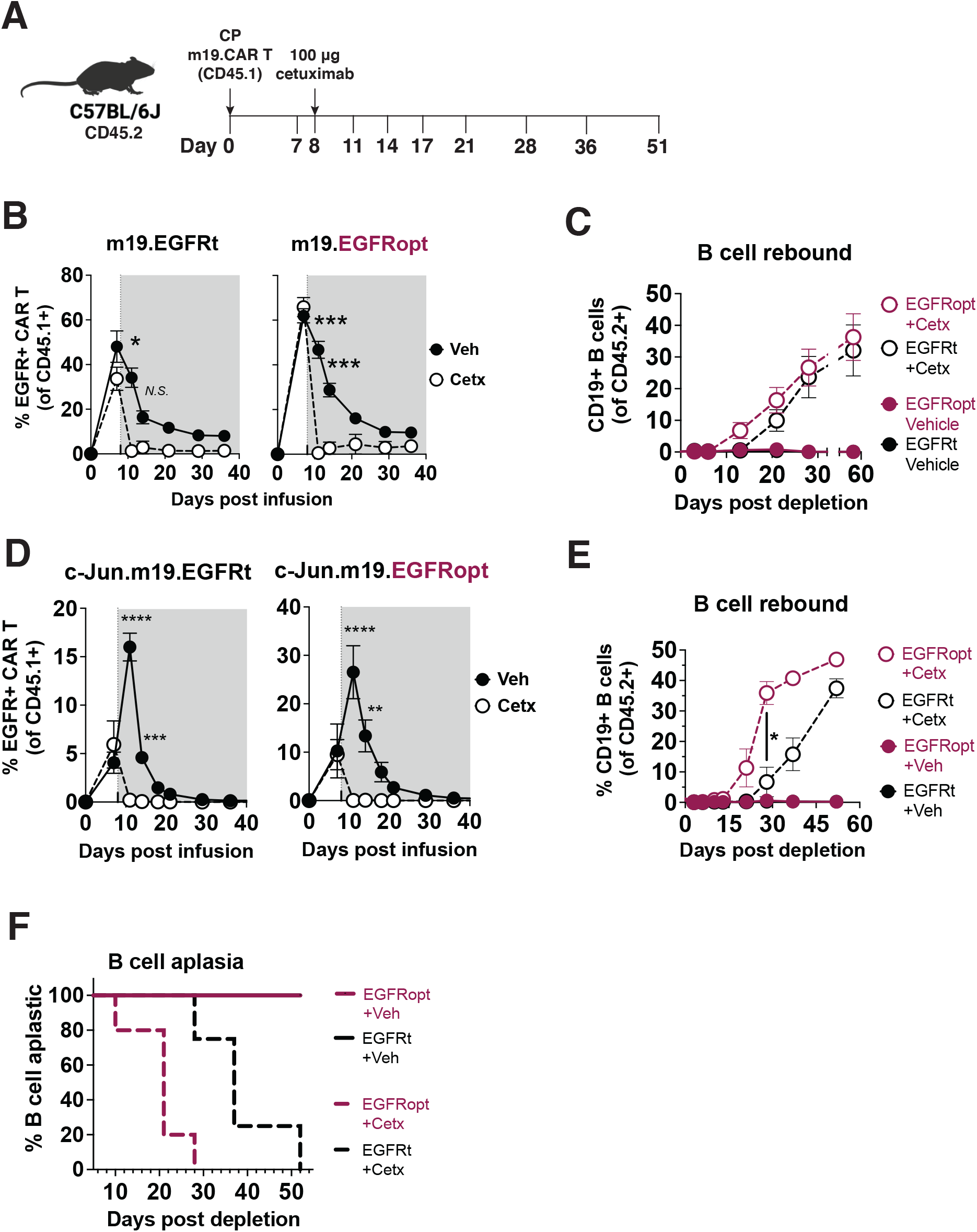
EGFRopt exhibits superior elimination of CAR T cell effector activity. (A) Schematic depicting the transfer and monitoring of the bicistronic m19.CAR T (B-C) or tricistronic c-Jun.m19.CAR T cells (D-F). (B) Frequencies of bicistronic m19.EGFRt and m19.EGFRopt CAR T cells following treatment with cetuximab (Cetx; white circle) or vehicle (Veh; black circle); the shaded area represents post depletion. (C) Kinetics of CD19+ B cell recovery in m19.CAR T cell-treated mice that received cetuximab or were left untreated. (D) Frequencies of tricistronic c-Jun.m19.EGFRt and c-Jun.m19.EGFRopt CAR T cells following treatment with cetuximab (Cetx; white circle) or vehicle (Veh; black circle); the shaded area represents post depletion. (E) Kinetics of CD19+ B cell recovery in c-Jun.m19.CAR T cell-treated mice that received cetuximab or were left untreated. (F) Frequencies of B cell-aplastic mice (containing < 3% CD19+ B cells of total live CD45+ cells) post cetuximab depletion of tricistronic c-Jun.m19.CAR T cells. (B-C) n=5-6 mice/group, (D-F) n=4-8 mice/group; symbols and error represent means ± SEM, with *P<0.05, **P<0.01, ***P<0.001, ****P<0.0001. Absence of error bar indicates SEM less than area represented by symbol. Analyzed by ANOVA (Šídák post hoc).

## DISCUSSION

Here we report the design of an optimized cell surface truncated EGFR safety switch that can be used to track and swiftly ablate transferred T cells *in vivo*. EGFRopt is a truncated, fully human sequence including a short charged juxtamembrane domain but lacking the ligand binding and receptor tyrosine kinase signaling domains. We demonstrate that the addition of three juxtamembrane arginine residues markedly increases the expression of truncated EGFR and enhances elimination of engineered T cells *in vitro* and *in vivo* using a commercially available antibody. EGFRopt is functionally inert; deletion of extracellular domains I and II as well as the intracellular kinase domain prevents binding and signal transmission by the native ligand of EGFR.^15^ Despite the removal of these domains, truncated EGFR retains its type I protein conformation and the epitope targeted by cetuximab. The remaining extracellular domains III and IV are stably expressed and do not have any known ligand or cell adhesion roles. Consistent with this, clinical testing of cell therapy constructs with surface expression of EGFR extracellular domains III and IV is safe, with no reported toxicity associated with expression of the EGFR-derived component.^19,31^ EGFRopt is a compact (~1kb), modular protein which can be easily integrated into next-gen cell therapy platforms as a single coding region or as part of a multi-gene cassette separated by 2A linkers.

The EGFR-based safety switch has advantages over previously described cell surface markers that offer dual use as transduction markers and safety switches. Studies utilizing RQR8, which incorporates a CD34 epitope and two CD20 mimotypes, demonstrated complete depletion of T cells expressing monocistronic RQR8 following a multi-dose regimen of rituximab^16^. However, when CAR T cells expressed bicistronic RQR8, depletion was incomplete due to downregulation of RQR8 post infusion.^32^ Further, truncated LNGFR^33^, CD19^14^, or CD34^34^ are expressed on endogenous immune cells and lack suitable pharmaceutical-grade cytotoxicity-inducing antibodies. By contrast, EGFR is not expressed in cells of the hematopoietic lineage and can be targeted by the commercially available, FDA-approved cetuximab (Erbitux). Our studies show that optimal targeting of EGFRt requires sufficient cell surface expression levels and that suitable short intracellular domains boost cell surface display of EGFR-derived proteins by 3-5 folds, likely by enhancing membrane insertion during protein synthesis or improving membrane protein stability. This enhanced surface expression of EGFRopt is also observed with large tricistronic constructs.

Targeting EGFRopt with cetuximab provided for more rapid and complete depletion of CAR T cells *in vivo*, including low CAR expressing T cells. Previous studies by ourselves^23^ and others^35,36^ demonstrated the utility of EGFRt as a safety switch for CAR T cells *in vitro* and in immunocompetent and xenograft models. However, only bicistronic constructs were used and cetuximab-mediated clearance of CAR T cells was slow, incomplete, and required multiple doses of cetuximab. The utility of EGFRt in multicistronic uses also remained unclear. For example, when blood-circulating EGFRt-marked CAR T cells were targeted with a high dose (1 mg) of cetuximab, they decreased by two-fold and required an additional week to decrease another two-fold after a second dose.^35^ Notably, cetuximab failed to significantly reduce the CAR copy in the bone marrow.^35^ Moreover, a study using monocistronic EGFRt+ cells in xenograft models demonstrated a two-log reduction of vector copy numbers in the bone marrow when 12 daily doses of 1mg cetuximab were administered.^36^ When bicistronic EGFRt cells were used, copy numbers decreased only by two-to-three folds in these tissues.^36^ Using EGFRopt+ CAR T cells, our studies revealed substantially more profound depletion in the blood 3 hours post a single low-dose of cetuximab and a nine-fold reduction in the bone marrow 24 hours later, compared to a mere two-fold reduction when EGFRt+ CAR T cells were targeted. Therefore, our work supports the conclusion that robust cell surface expression of truncated EGFR elicits swift and complete cetuximab-mediated ADCC. This enhanced EGFRopt expression does not influence the CAR T cell cytotoxic potential, as both EGFRt+ and EGFRopt+ CAR T cells mediated B cell aplasia for the duration of study.

Our studies are limited to characterizing EGFRopt in T cells and do not address its use for ablating engineered innate immune cells such as macrophages and NK cells. Cetuximab poses certain clinical limitations with therapeutic dosing including adverse reactions in 80% of patients, with 9-19% developing III/IV adverse events^37^ including acneiform follicular skin exanthema, abdominal pain, nausea, and asthenia.^38^ The dose and frequency of cetuximab used for targeting EGFRopt would be expected to elicit fewer adverse events than those associated with the high-dose treatment of EGFR-positive malignancies. While the fully human IgG2 panitumumab targets the same epitope as the chimeric IgG1 cetuximab but does not elicit toxicity,^39^ the current studies did not address whether panitumumab can induce sufficient ADCC upon binding EGFRopt *in vivo*. Lastly, our *in vivo* observations of the EGFRopt safety switch are also limited to immunocompetent murine models, and the safety and efficacy of EGFRopt in patients remains to be investigated.

In summary, this work illustrates the utility of EGFRopt as a superior safety switch for next-generation, genetically engineered T cell therapies. The more complete and swift depletion kinetics of infused cells expressing EGFRopt may increase its utility for managing acute toxicities resulting from infused T cells. As targets for cell therapy expand and new technologies enhance cell persistence and function, strategies for cell ablation which use approved clinical reagents will become an increasingly important feature.

## MATERIALS AND METHODS

### Viral constructs

Lentiviral vectors under the control of MND promoter included the following coding sequences in order: ROR1-specific CAR derived from R12 scFv with CD28 transmembrane domain, 4-1BB and CD3z signaling domains, a P2A self-cleaving peptide, and truncated EGFR including domains III, IV, and transmembrane domain (EGFRt)^40^ or an EGFR variant incorporating an additional juxtamembrane domain. Constructs with tricistronic cassettes were linked in frame, placed under the control of MND, and included: human c-Jun, P2A peptide, ROR1-specific CAR, P2A peptide, and EGFRt or an EGFR variant incorporating an additional juxtamembrane domain. The R12 scFv was derived from the R12 anti-ROR1 antibody.^41^

MP71 retroviral vectors were used to transduce murine T cells and encoded: murine CD8α signal peptide (Uniprot P01731 aa1-27), murine-CD19-specific scFv (hereafter ‘m19’), murine CD8α hinge and transmembrane (Uniprot P01731 aa151-219), murine CD28 (Uniprot P31041 aa177-218), and murine CD3z (Uniprot P24161 aa52-164), linked by a P2A peptide sequence to mouse-codon-optimized truncated human EGFR (EGFRt; Uniprot P00533 aa334-668) or EGFRopt (aa334-671). Where indicated, Strep Tag II (NWSHPQFEK) and (G4S)2 linked the scFv to hinge. To generate tricistronic constructs, murine c-Jun (Uniprot P05627 aa1-334) was cloned upstream of m19.CAR, linked by a T2A sequence.

### Retrovirus production

Lentivirus was produced by transient transduction of transfer plasmid and 3rd generation helper plasmids using polyethylenimine (Polyplus) into HEK 293T cells (ATCC). Lentivirus containing supernatants were collected 3 days later, filtered through a 0.22 μm filter, concentrated overnight with Lenti-X (Takara), and frozen at −80°C. Lentivirus was titered prior to use. For retrovirus production, Plat-E cells (Cell Biolabs) were transiently transfected using calcium phosphate (Takara) and retrovirus-containing supernatants were collected 2 days later, filtered through 0.45 μm filter, snap frozen on dry ice prior for storage, and later titered.

### Human T cell culture and viral transduction

Pre-selected, cryopreserved primary human CD4+ and CD8+ T cells from normal donors were obtained from BloodWorks (Seattle, WA). T cells were cultured in OpTmizer medium (ThermoFisher) supplemented with Immune Cell Serum Replacement (ThermoFisher), 2 mM L-glutamine (Gibco), 2 mM Glutamax (Gibco), 200 IU/ml IL-2 (R&D systems), 120 IU/mL IL-7 (R&D systems), and 20 IU/mL IL-15 (R&D systems). For lentiviral transduction, the T cells were stimulated with 1:100 dilution of T cell TransAct (Miltenyi) for 30 hours. Pre-titered virus was then added to the T cells for 18-24 hours. Stimulation and viral infection were terminated by addition of 7 volumes of fresh media and cytokines without TransAct, and cells were cultured for 3-7 additional days before analysis.

### Murine T cell viral transduction and adoptive transfer

CD8 T cells were negatively selected (Stem Cell) from spleen and peripheral lymph nodes of 6-8 week-old CD45.1 mice, and stimulated with 1 mg/mL plate-bound anti-CD3 (145-2C11) and anti-CD28 (37.51) for 20 hours at 37°C 5% CO2 in complete RPMI (RPMI-1640, 10% heat-inactivated FBS, 1 mM HEPES, 100 U/mL penicillin/streptomycin, 1mM sodium pyruvate, and 50 *μ*M b-mercaptoethanol) supplemented with 50 U/mL murine IL-2 (Peprotech). Retrovirus was centrifuged for 2 hrs at 2560rcf at 32^0^C onto wells pre-coated with RetroNectin (Takara). Wells were rinsed with PBS and CD8 T cells were added at 1×10^6^ cells/mL in complete RPMI supplemented with 50 U/mL IL-2 and mouse T-activator Dynabeads (ThermoFisher) at 1:1 ratio. Plates were centrifuged at 800rcf for 30 mins at 32^0^C and incubated overnight. IL-2-supplemented complete RPMI media was replaced, and T cells were incubated for an additional 24 hrs. T cells were harvested, resuspended at 1×10^6^ cells/mL in complete RPMI supplemented with 50 U/mL murine IL-15 (Peprotech) and incubated for additional 48 hrs. Activator beads were removed and transduction efficiency was determined by flow cytometry. CAR T cells were prepared for infusion by resuspending at 3×10^6^ EGFR+ cells per 100 *μ*L serum-free RPMI-1640 and kept on ice prior to adoptive transfer. 6-8 week-old C57BL/6J female mice were pre-conditioned with intraperitoneal injection of 200 mg/kg cyclophosphamide and 6 hr later were injected intravenously with 3×10^6^ CAR T cells.

### Flow Cytometry

For analysis of human T cells, 1-2×10^5^ cells were stained using Fixable Viability Dye eFluor780 (eBioscience), and antibodies against EGFR-A488 (Hu1 cetuximab biosimilar; R&D), EGFR-BV421 (AY13; Biolegend), CD3-BUV805 (SK7, ThermoFisher), and CD56-PE (HCF56; Biolegend) on ice for 30min. For detecting R12 CAR, purified recombinant ROR1-Fc was produced in-house and conjugated to Alexa647. Cells were washed twice with staining buffer (eBioscience) before and after staining, fixed in FluoroFix Buffer (BioLegend), stored at 4°C in the dark until acquisition on ZE5 cytometer (Bio-Rad). For analysis of mouse samples, 50 *μ*L peripheral blood was collected into EDTA-coated tubes and underwent two rounds of ACK lysis prior to surface staining. Samples were stained using Live/Dead Fixable Aqua Dead Cell stain (Invitrogen) at 4^0^C for 15min. Cells were stained for 30min in flow buffer (PBS, 1mM EDTA, 2% FBS) with antibodies against CD8α-FITC (53-6.7; Biolegend unless otherwise noted), CD19-PerCP/Cy5.5 (1D3), CD4-PE-Cy7 (GK1.5), CD45.2-APC/Cy7 (104), CD45.1-BrV421 (A20), hEGFR-APC (AY13) and acquired on BD Celesta. Cells were alternately stained with antibodies against CD8α-FITC APC (53-6.7), hEGFR-PE (AY13), and biotinylated anti-STII (5A9F9, Genscript), followed by SAv-DyLight488. Data analysis was performed on FlowJo 10 software (Treestar).

### Antibody dependent cellular cytotoxicity

Natural killer cells were isolated from cryopreserved, T cell depleted (CD4-/CD8-) PBMC (AllCells) by negative selection using the EasySep Human NK Cell Kit (Stemcell) according to the manufacturer’s protocol. To activate their cytolytic function, isolated NK cells were cultured in RPMI-10 supplemented with 10 ng/ml human IL-15 overnight before use.^42,43^ Cryopreserved, transduced primary T cells were thawed and pre-cultured overnight in OpTmizer medium plus cytokines as described above. The cells were then counted, resuspended in RPMI-10, and added to a V bottom 96 well plate in a 100 *μ*l volume and incubated with a cetuximab biosimilar at the indicated final concentration. IL-15 primed NK cells were then added at a 10:1 ratio of NK:CAR-T cells and the V bottom plate was gently centrifuged (100xg, 30 sec) to bring effector and target cells together. After co-culture, remaining CAR+ T cells were identified by FACS. Samples were stained with anti-CD3, anti-CD56, ROR1-Fc, and FVD780, fixed, and acquired under volumetric counting mode. Antibody specific ADCC of T cells was assessed by comparing the total live CD56-CD3+ROR1 -Fc+ populations in conditions treated or not treated with the antibody.

### In vivo cetuximab treatment

For depletion of transferred EGFRt+ or EGFRopt+ CAR T cells, cetuximab (Lilly) was infused at the indicated doses on Day 8 or 14. Mice were determined to exhibit B-cell aplasia when CD19+ B cell frequency in peripheral blood declined to below 3% of the total circulating endogenous CD45.2+ cells as determined by flow cytometry.

### Gene expression analysis

Red-blood-cell lysis using ACK buffer was performed on 25-35 μL peripheral blood, total RNA was isolated using RNeasy micro kit (Qiagen), and complementary DNA (cDNA) was amplified using SuperScript IV First-Strand Synthesis System (Invitrogen). Gene expression was measured by qRT-PCR on ABI QuantStudio 5 using TaqMan Universal Master Mix II with UNG (Applied Biosystems) with pre-validated primers for murine *b2m* (IDT PrimeTime assay Mm.PT.39a.22214835; Ref NM_009735) or custom-designed primers for *egfr* transgene (IDT PrimeQuest Tool; FWD CCAACACCATCAACTGGAAGA; REV GCTACACAGAGCGTGACAAA; Probe TCATCAGCAACAGAGGCGAGAACA). Gene expression was measured as 2-(Δ*C*_⊤_), where Δ*C*_⊤_=*C*_⊤_*egfr–C*_⊤_*b2m*, and fold changes were then calculated as 2-^(Δ*C*_⊤_relerence sample-_⊤_tested sample)^.

### Study approval

Animal studies were performed in accordance with institutional IACUC rules (protocol No.50884). C57BL/6J (CD45.2) and B6.CD45.1 donor mice were acquired from Jackson Laboratory and housed at the FHCC.

### Statistics

Statistical differences were determine using ratio paired t test or ANOVA with Šídák post-test. Mice were randomized when assigned to treatment groups. Statistical analyses were performed using Prism 9 (GraphPad), and differences were considered significant if P ≤ 0.05.

## ACKNOWLEDGMENTS

The authors thank D. Parilla, L. King, and E. Gad for their assistance in performing mouse studies; A. Chen for input on qRT-PCR primer custom design; C. Simianer and S. Spadinger for preparing T cells used in ADCC assays; C. Wu for lentiviral production; and S. Boyken, B. Sather, S. Park, Q. Vong, and B. Boldajipour for helpful discussion. This work was supported by funding from Lyell Immunopharma, Inc. The MP71 retroviral vector was a gift from W. Uckert (Max Delbruck Center for Molecular Medicine). S.R.R. is a co-founder of Lyell Immunopharma, Inc. and has received grant funding from Lyell Immunopharma, Inc. S.R.R. was a co-founder Juno Therapeutics, a Bristol Myers Squibb company and has served as a scientific advisor to Juno Therapeutics and Adaptive Biotechnologies. S.P. is an employee of Lyell Immunopharma, Inc. M.J.L. is a co-founder of Lyell Immunopharma, Inc., and Outpace Bio. H.F.M. and M.J.L. are listed as inventors on patent applications related to this work. H.F.M., L.T., and M.J.L. are employees of Outpace Bio, a licensee of patent rights owned by Lyell Immunopharma, Inc. The remaining authors declare no competing interests.

## AUTHOR CONTRIBUTIONS

T.B.S., H.F.M., S.M.S., T.D., S.P., and L.J.T. acquired data and performed statistical analysis. T.B.S., H.F.M., M.J.L and S.R.R. conceived and designed research. T.B.S., H.F.M., M.J.L., and S.R.R wrote the manuscript.

