## Supplemental Figure 1 for "Safety switch optimization enhances antibody-mediated elimination of CAR T cells"

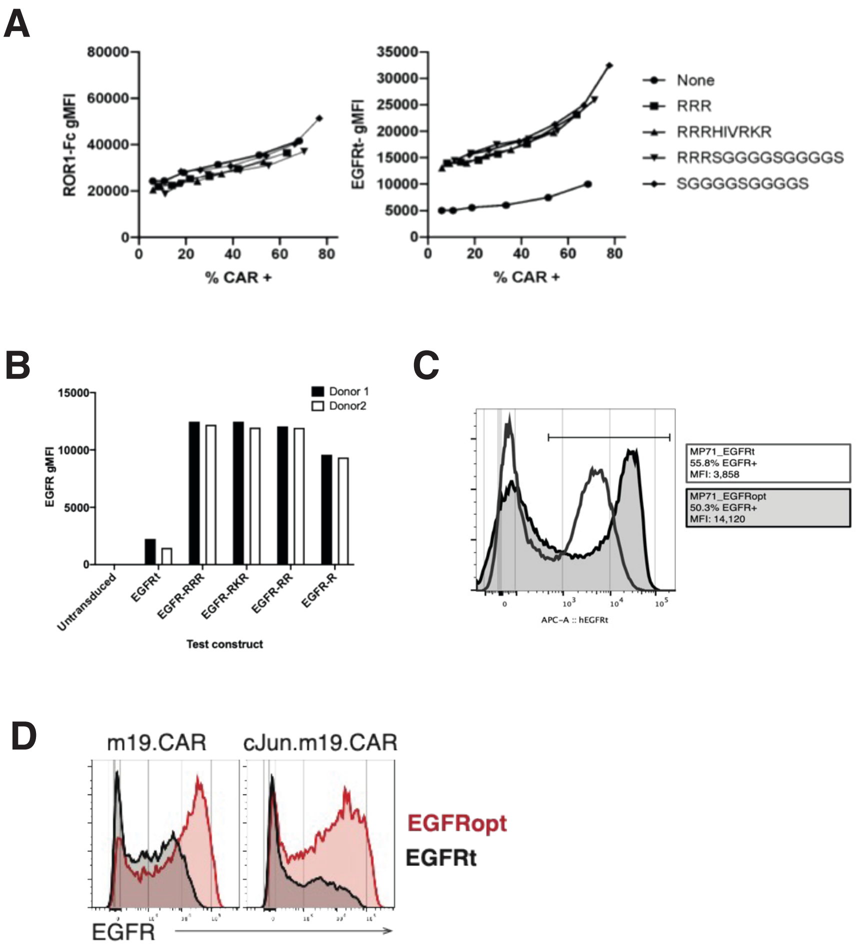


**Supplementary Figure 1: Surface expression of EGFR-derived proteins**

(A) Juxtamembrane domains improve surface display of EGFR across a wide range of transduction (left) without affecting surface CAR-expression (right) on CAR+ transduced T cells. (B) gMFI values of truncated EGFR molecules that include no juxtamembrane sequence (EGFRt), one arginine residue (R), two arginine residues (RR), three arginine residues (RRR), or one arginine residue swapped for lysine (RKR) encoded in a tricistronic construct. (C) Expression levels of monocistronic EGFRt (white) and EGFRopt (grey) on murine CD8$^{+}$ T cells with similar transduction efficiencies. (D) Expression levels of EGFRt and EGFRopt on murine CD8$^{+}$ T cells transduced with the bicistronic (left) and tricistronic (right) constructs.
